# p53 suppresses mutagenic RAD52 and POLθ pathways by orchestrating DNA replication restart homeostasis

**DOI:** 10.1101/204057

**Authors:** Sunetra Roy, Karl-Heinz Tomaszowski, Jessica Luzwick, Soyoung Park, Jun Li, Maureen Murphy, Katharina Schlacher

**Affiliations:** Department of Cancer Biology, UT MD Anderson Cancer Center, Houston, TX, USA.; Department of Genomic Medicine, UT MD Anderson Cancer Center, Houston, TX, USA; Molecular and Cellular Oncogenesis Program, The Wistar Institute, Philadelphia, PA, USA.

**Keywords:** p53, genome instability, cancer, replication fidelity, replication restart, fork reversal, fork protection, MLL3, MRE11, RAD52, POLq, cancer drug resistance, Fanconi Anemia, BRCA1, BRCA2

## Abstract

Classically, p53 tumor-suppressor acts in transcription, apoptosis, and cell-cycle arrest. Yet, replication-mediated genomic instability is integral to oncogenesis, and p53 mutations promote tumor progression and drug-resistance. By delineating human and murine separation-of-function p53 alleles, we find that p53 null and gain-of-function (GOF) mutations exhibit defects in restart of stalled or damaged DNA replication forks driving genomic instability independent of transcription activation. By assaying protein-DNA fork interactions in single cells, we unveil a p53-MLL3-enabled recruitment of MRE11 DNA replication restart nuclease. Importantly, p53 defects or depletion unexpectedly allow mutagenic RAD52 and POLθ pathways to hijack stalled forks, which we find reflected in p53 defective breast-cancer patient COSMIC mutational signatures. These data uncover p53 as a keystone regulator of replication homeostasis within a DNA restart network. Mechanistically, this has important implications for development of resistance in cancer therapy. Combined, these results define an unexpected role for p53 suppression of replication genome instability.

## Introduction

One of the most prominent hallmarks of cancer is genomic instability (Hanahan and Weinberg, 2011). As such, many DNA damage response or repair genes that restore genome stability are known tumor suppressors, including p53, the guardian of the genome (Kim et al., 2015). In breast cancers, p53 mutations are associated with more aggressive and triple negative breast cancers (Turner et al., 2013). Similar to high serous ovarian cancers, these aggressive cancers respond to chemotherapy including platinum drugs and PARP inhibitors initially, but develop resistance thereafter (Luvero et al., 2014; Wahba and El-Hadaad, 2015).

First thought to be a proto-oncogene, the initial discovery of a gain-of-function (GOF) p53 mutant allele (Lane and Crawford, 1979; Levine and Oren, 2009; Linzer and Levine, 1979) masked the loss of wild-type (WT) p53 function. Despite early discrepancies, only a decade later p53 was recognized as a tumor suppressor (Baker et al., 1989). Loss of p53 function can occur either by deletion or by mutation that otherwise may exhibit a GOF, typically enhancing transcription functions. To date, the most consistent defect for both null and GOF p53 mutants in cancers relates to p53’s transcription factor function to promoting apoptosis and cell cycle arrest.

Genetic data of several separation-of-function p53 mutant mice suggest that there are additional p53 functions that contribute to tumor progression, which are transcription independent; Murine p53 H172P corresponding to human H175P retains much if its tumor suppression function despite loss of transcriptional induction of p21 and loss of apoptosis (Liu et al., 2004). Similarly, p53 mutations in the transactivation domain and p53 acetylation mutations severely inhibit p53 apoptosis and senescence, yet exhibit a mild and delayed tumor onset (Li et al., 2012; Zhu et al., 2015). p53 also has seemingly disparate cellular functions including during metabolism and epigenetic control, i.e. through its interaction with MLL3/4 histone methyltransferases (Pfister et al., 2015; Zhu et al., 2015), although the contribution of these functions to tumor suppression is not fully understood.

For cancer, a prominent p53 function is to maintain genomic stability upon DNA damage as part of a damage response. Since DNA damage traditionally is most prominently considered in the context of double-strand break (DSB) lesions, many studies focus on putative p53 functions in DSB repair. Next to error-free repair of DSBs by homologous recombination (HR) involving BRCA1/2 and RAD51, DSBs may also be repaired by non-homologous end joining (NHEJ), or through secondary and typically mutagenic pathways of single strand annealing (SSA) mediated by RAD52 and micro-homology mediated end joining (MMEJ) involving POLθ (Black et al., 2016; Branzei and Foiani, 2008; Moynahan and Jasin, 2010; Wood and Doublie, 2016). In response to DNA damage, ATM mediates p53 phosphorylation as part of a DNA stress response (Saito et al., 2002), which is facilitated by PTIP (Jowsey et al., 2004), a BRCT domain containing protein that is part of the MLL3/4 complex.

Similar to its transcription function, discrepancies ensue in the molecular function of p53 during DNA repair. While indirect studies found p53 to inhibit error-free homologous recombination (HR) and spontaneous sister-chromatid exchange (SCE), which somewhat paradoxically was proposed to promote genomic stability (Bertrand et al., 2004; Gatz and Wiesmuller, 2006), loss of p53 does not change DSB repair rates by HR when measured in specific induced break assays (Willers et al., 2001). Thus, the mechanism by which p53 promotes genomic stability associated with tumorigenesis remains contradictory.

As previously hypothesized (Cox et al., 2000), recent findings formalized that 2/3 of all mutations found across cancers are caused by errors occurring during proliferation (Tomasetti et al., 2017), highlighting the critical importance of protective mechanisms during DNA replication. Intriguingly, early studies found p53 is activated at stalled replication forks (Gottifredi et al., 2001; Kumari et al., 2004), which are a source for genomic instability requiring distinct replication fork stability pathways (Branzei and Foiani, 2010). p53 interacts with BLM helicase at replication forks and represses HR in S-phase upon DNA damage, independent of its G1-S and transactivation activity (Bertrand et al., 2004; Janz and Wiesmuller, 2002; Saintigny and Lopez, 2002). Hinting at a direct p53 replication function, recently p53 was found to interact with DNA POL*i* (Hampp et al., 2016). Moreover, p53 deletion in U2OS cells was reported to slow unperturbed replication (Klusmann et al., 2016), although this was suggested to require p53 transcription function, while p53 functions in DNA damage response are not.

Here we identify a critical role for p53 in balancing replication pathway homeostasis and show p53 suppresses replication genomic instability independent of transcription activation. We find p53 mutant alleles that separate transcription activation and replication restart functions and reveal a direct correlation between p53 replication and tumor progression functions. Importantly, we find mutagenic RAD52/POLθ replication pathways increase for both GOF and p53 null alleles in a transcription independent manner, consistent with mutation signatures that we identify in p53 mutant breast cancers. Our results thus allow for an unexpected alternative hypothesis for acquisition of resistance in breast cancer cells due to p53 loss: mutant p53 boosts mutagenic RAD52/POLθ pathways, which increase deletion and point mutations that can lead to secondary resistance mutations.

**eLife digest** p53 regulates replication restart pathway homeostasis by recruiting chromatin-remodeler MLL3 and restart-nuclease MRE11. It so suppresses binding of mutagenic RAD52/POLθ to DNA replication forks, which is independent of its transcription activation function. The combined data defines a role for p53 in suppressing genome instability during replication.

## Results

### Transcription-independent p53 function for restart of stalled replication forks

p53 is an ATM phosphorylation target, is activated at stalled replication forks (Kumari et al., 2004) and interacts with BLM helicase. As BLM helicase is implicated in replication restart (Davies et al., 2007), as is ATM (Trenz et al., 2006), we tested the role of p53 in DNA replication reactions when stalled with dNTP depleting hydroxyurea (HU). Using single-molecule DNA fiber spreading (Figure 1A), we assessed the number of stalled replication forks after low-dose replication stalling (Figure 1A), as a test for defects in replication restart. We find a doubling of stalled forks in CRISPR/CAS9-engineered p53-null human HAP-1 cells compared to cells with wild-type (WT) p53 (Figure 1B; 35% stalled forks in p53 null 18% WT p53 HAP-1). This suggests a prominent role for p53 in the resumption of DNA replication after replication stress.

**Figure 1 with 1 supplement.**
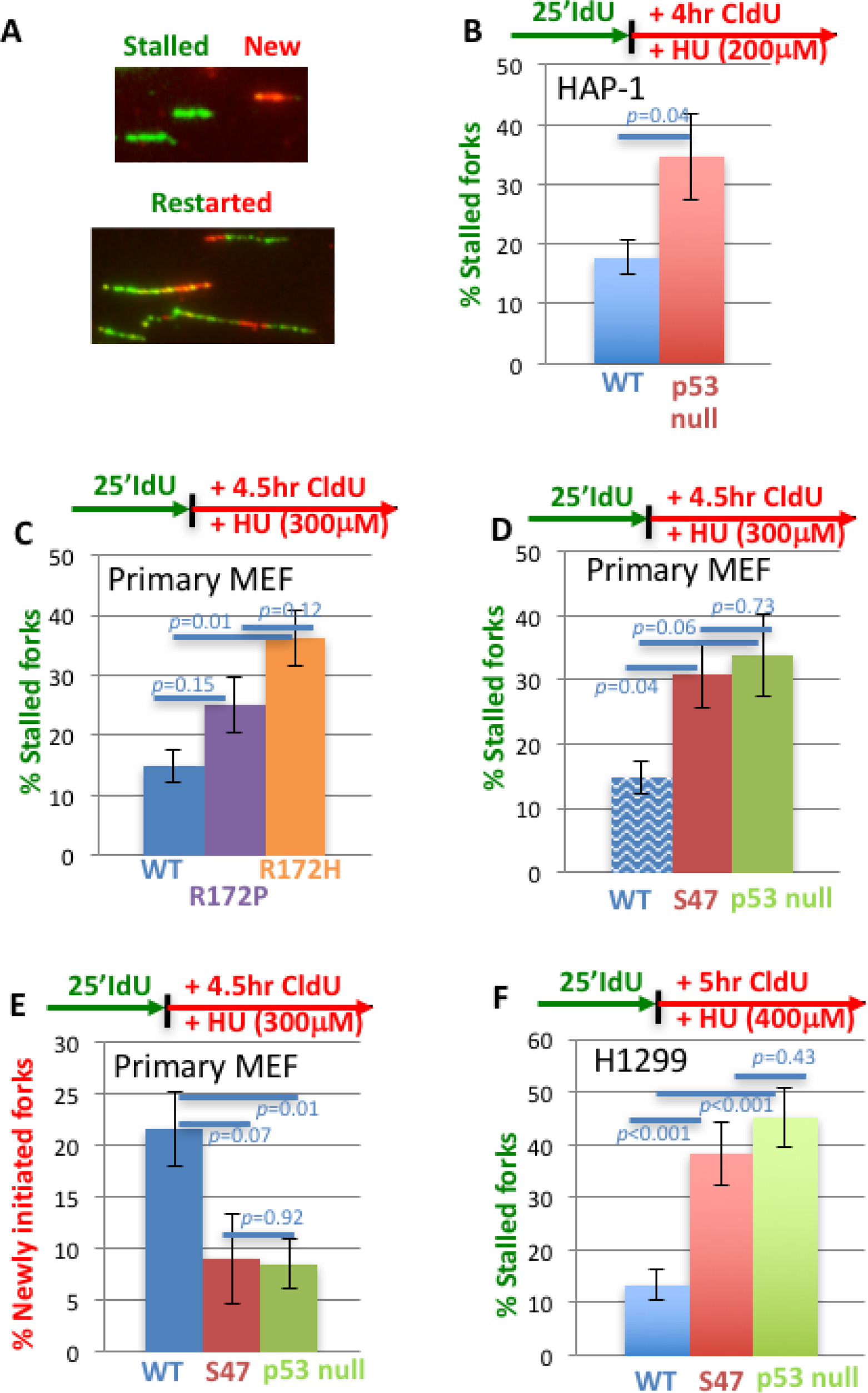
p53 promotes efficient replication restart of stalled DNA forks. (**A**) Representative image of DNA fibers. The number of stalled forks (# stalled/(# stalled +#restarted) or newly initiated origins ((# CldU/(# stalled +# restarted +#new) is obtained in the following panels.(**B**) HAP-1 p53 null cells (**C**) primary MEF with GOF p53 R172P (apoptosis deficient, mildly tumor prone) and p53 R172H (apoptosis deficient, strongly tumor prone), (**D,E**) primary MEF with p53 S47 (apoptosis and greatly transcription proficient) and null for p53 (for better comparison WT p53 of (C) is re-plotted, shady blue), and (**F**) human H1299 cells with inducible p53 WT or S47. Error bars represent the SEM. Significance values are derived from student T-test analysis.

Increased fork stalling is classically compensated for by increased new origin firing, as seen for CHK1 defects (Petermann and Helleday, 2010; Petermann et al., 2010). Unexpectedly, we find that increased fork stalling in p53 null cells is accompanied by a decrease, rather than an increase, in new origin firing compared to both WT p53 (Figure 1 – figure supplement 1A; 20% newly fired origins in HAP-1 cells, respectively, and 9% newly fired origins in HAP-1 p53 null cells). Taken together, the data suggests that p53 defects in HAP-1 cells exhibit distinct and unconventional replication restart defects resulting in both decreased replication restart and decreased new origin firing.

In tumors, p53 is deleted or mutated typically resulting in gain-of-function (GOF) (Freed-Pastor and Prives, 2012). To test whether p53 mutations alter replication restart, we investigated one of the most common GOF mutations in primary mouse embryonic fibroblasts (murine p53 R172H corresponding to human p53 R175H) (Liu et al., 2000). These cells show both an increase in stalled replication forks, and fewer restarted forks compared to WT p53 MEFs (Figure. 1C; 36% stalled forks in p53 R172H MEF and 15% in WT MEF and Figure 1 – figure supplement 1B). These data thus uncover conserved defective outcomes for both null and GOF p53 mutations at stalled replication forks.

Murine p53 R172P corresponds to a rare human polymorphism R175P, which results in loss of transcriptional activation and apoptosis resembling p53 null (Liu et al., 2004). Yet, tumor development is markedly less severe in p53 R172P mice compared to p53 null mice, suggesting alternative mechanisms contribute to tumor progression besides transcriptional regulation of apoptosis with this mutation (Liu et al., 2004). We therefore tested p53 R172P MEFs and find that they only show a moderate increase in fork stalling compared to p53 R172H (Figure 1B). Thus, restart functions in p53 R172 mutant cells show an improved correlation with tumor suppression activity *in vivo*.

To further examine the possible link of failed p53 replication restart function and cancer, we tested p53 S47 (P47S), which is a breast-cancer pre-disposition polymorphism in African-descent populations (Jennis et al., 2016). It is largely transcriptionally active including for p21 (Felley-Bosco et al., 1993; Jennis et al., 2016); it remains proficient for apoptosis (Felley-Bosco et al., 1993; Jennis et al., 2016). Yet, p53 S47 mice are tumor prone, and p53 S47 contributes to breast cancer risk in African populations (Jennis et al., 2016; Murphy et al., 2017). We find that S47 slows cell growth similar to WT p53 (Figure 1 - figure supplement 1C) consistent with intact cell-cycle check-point functions. Furthermore, we find that p53 S47 primary MEFs exhibit a loss-of-function (LOF) for replication restart, as measured by an increase in stalled forks upon HU treatment compared to WT MEF cultures (Figure 1D; 31% stalled forks in p53 S47 MEF versus 15% in WT MEF). Moreover, p53 S47 MEFs resemble p53 null MEFs in their inability to restart forks (Figure 1D; 34% stalled forks in p53 null MEF). Similar to p53 null HAP-1 and GOF R172H MEF cells, both p53 null MEF and p53 S47 MEFs show defects in new fork initiation (Figure 1E; 21% newly fired origins in p53 WT MEF versus 9% and 8% in p53 S47 MEF and p53 null MEF).

To test whether p53 S47 restart defects are conserved in human cells, we expressed human p53 S47 under doxycycline control in H1299 non-small cell lung carcinoma cells (Figure 1 - figure supplement 1D) and examined stalled forks. We found an increase in stalled forks that resembles p53-null H1299 cells. Both p53 null and p53 S47 H1299 cells exhibit a substantial increase in stalled forks compared to WT p53 expressing H1299 cells (Figure 1F; 13% stalled forks in p53 WT, 38% in p53 S47, and 45% in null H1299 cells). Taken together, these results suggest that p53 promoted replication restart is largely independent of its transcription-activation function in cell cycle progression and apoptosis.

### p53 restart defects promote replication-dependent genome instability and cellular sensitivity to replication stalling agents

Restart defects cause cellular sensitivity to replication stalling agents, as seen for cells with BLM defects (Davies et al., 2004). We reasoned that in mutant p53 cells, this cellular phenotype so far may have been obscured by loss of apoptosis, which can override cellular sensitivity by inhibiting cell death. We therefore tested cellular sensitivity using the loss-of-function (LOF) mutant p53 S47, which remains largely apoptosis proficient. We find that p53 S47 expressing H1299 cells are sensitive to replication stalling agents HU (Figure 2A) and mitomycin C (MMC; Figure 2B). This cellular replication stress phenotype is masked when the p53 mutations additionally inactivate apoptosis and cell-cycle check-point functions, such as in p53 null H1299 cells (Figures 2A and 2B) or p53 null compared to WT mammary epithelial MCF10A cells (Figure 2 - figure supplement 1). Collectively, the data with apoptosis-proficient p53 S47 implies that p53 functions in replication restart suppresses cellular sensitivity to replication stress.

**Figure 2 with 1 supplement.**
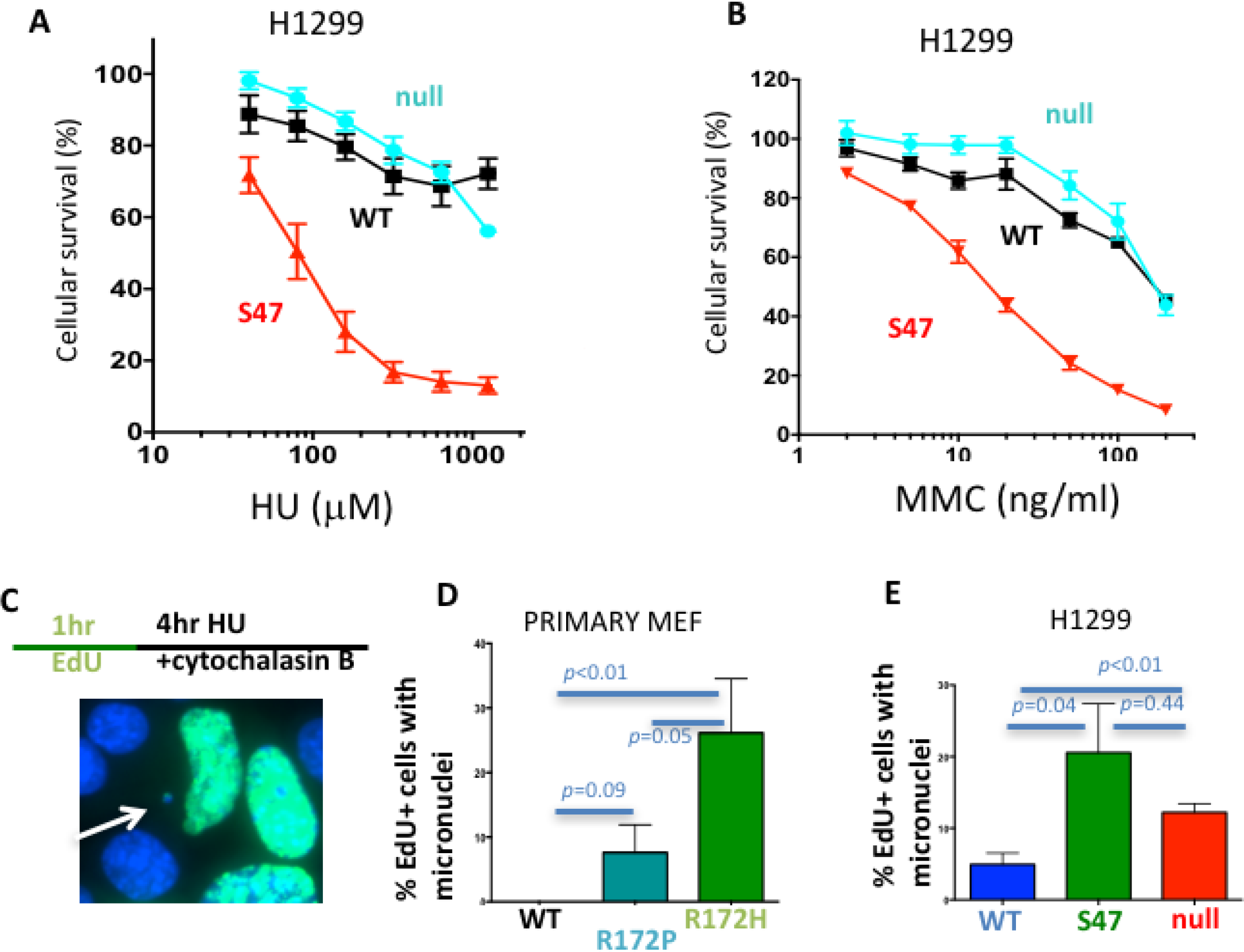
p53 promotes replication dependent genomic stability. (**A**) Cellular sensitivity to replication stalling with hydroxyurea (HU) in H1299 cells (**B**) Cellular sensitivity to replication stalling with mitomycin C (MMC) in H1299 cells (**C**) Experimental scheme and representative image of micronuclei; Scored are EdU positive cells with micronuclei only that were in S-phase during replication stalling with HU and that are blocked during cytokinesis immediately following the offending S-phase (**D**) Micronuclei in primary p53 R172P and R172H MEF and (**E**) human H1299 p53 WT, S47 and null cells. Error bars represent the SEM. Significance values are derived from student T-test analysis.

Separation-of-function mutations p53 R172P and S47 reveal a feasible correlation between loss of p53 restart function and tumor progression. We therefore tested whether p53’s replication restart function could contribute to genomic instability, which is a hallmark of cancer (Hanahan and Weinberg, 2011). Unresolved replication forks can result in DNA bridges, which convert to micronuclei, a mark of BLM defective cells (Hoffelder et al., 2004). We assessed genome instability by scoring micronuclei in p53 R172P, R172H and WT MEFs after arrest in cytokinesis immediately following replication stalling (Figure 2C). By considering EdU-positive cells only, the experimental set up ensures that only micronuclei are scored that result from induced replication stalling during the preceding S-phase. Consistent with an intermediate phenotype for replication restart, primary MEF p53 R172P exhibited less micronuclei with replication stalling compared to p53 R172H (Figure 2D; average of 8% micronuclei in p53 R172P and 26% in p53 R172H) albeit considerably more than WT p53 MEFs (none detected in WT).

With replication stalling, human p53 S47 expressing cells showed a marked increase in cells containing micronuclei compared to WT p53 expressing H1299 cells (Figure 2E average of 20% in p53 S47 and 5% in p53 WT H1299 cells). Micronuclei instability in p53 null H1299 cells similarly is significantly higher than in WT p53 H1299 cells (Figure 2E; average of 12%). Taken together, the genomic instability data corresponds with restart defects found in the respective p53 mutants, and by implication can contribute to tumor suppression.

### p53 promotes MLL3-chromatin remodeler and MRE11 restart nuclease recruitment to forks

p53 interacts with chromatin remodeling complexes and is implicated in facilitating epigenetic alterations (Pfister et al., 2015; Zhu et al., 2015). We observed an atypical restart defect in p53 mutant and null cells with less newly initiated replication forks (Figure 1E and supplement 1, 1A and 1B). We hypothesized that replication restart and new replication fork firing may require local chromatin opening and epigenetic alterations mediated by p53, which could explain the unusual decrease in new fork firing with p53 mutations. To investigate local protein changes specific to replication forks, we developed the SIRF assay (in Situ Interactions at Replication Forks using PLA; Figure 3A). Specifically, we applied sensitive proximity ligation chemistry to detect interactions between nascent, EdU labeled DNA and proteins within nanometer proximity. The signal is specific as elimination of EdU results in no signals (Figure 3A). MLL3 promotes H3K4 histone methylation to mark open chromatin (Ruthenburg et al., 2007). MLL3 association with replication forks in unperturbed cells is similar in p53 null and WT HAP-1 cells (Figure 3B;11 MLL3-bound replication sites per cell in WT p53 and 12 sites in p53 null HAP-1 cells). Upon replication stalling with HU, we see a marked increase in MLL3-bound replication sites in WT, but not in p53 null human HAP-1 cells (Figure 3C; 17 MLL3-bound sites in WT and 13 in p53 null HAP-1 cells). These data suggest inefficient MLL3 recruitment to forks upon replication stalling in the absence of p53.

**Figure 3 with 1 supplement.**
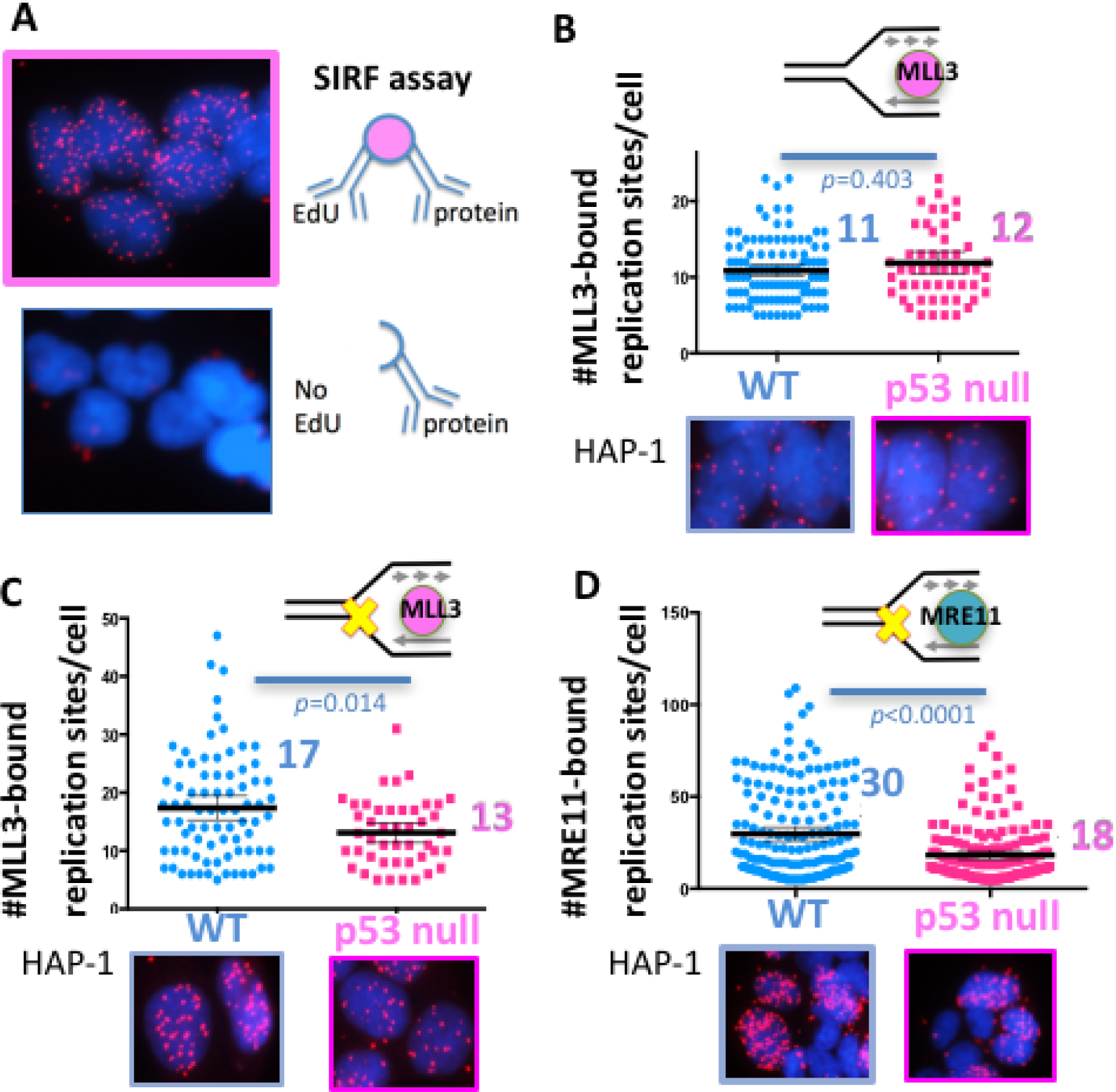
p53 promotes recruitment of chromatin remodeler and MRE11 to stalled replication forks. (**A**) Schematic and representative image of SIRF (In Situ Interactions at Replication Forks) assay for interactions between protein and nascent DNA in single cells: nascent DNA is pulse-labeled with EdU a protein of interest is crosslinked to the DNA immediately following the EdU pulse. Alternatively, EdU is washed out and cells are incubated with HU (200-400μM HU) before crosslinking. Proximity ligation assay (PLA) amplification with antibodies against EdU and the protein of interest will result in a signal only if interactions between the nascent DNA and the protein of interest are in close proximity. No signal is produced if the cell has not incorporated EdU. (**B**) Quantitation of SIRF assay of epigenetic remodeler MLL3 at unchallenged replication forks and (**C**) at HU stalled replication forks (yellow x) in HAP-1 p53 null and WT cells. (**D**) Quantitation of SIRF assay of MRE11 at HU stalled replication forks in HAP-1 p53 null and WT cells. Bars represent the mean and the 95% confidence interval. Significance values are derived from Mann-Whitney analysis.

MLL3-mediated chromatin opening is implicated in MRE11 nuclease recruitment to stalled replication forks (Ray Chaudhuri et al., 2016), a repair nuclease that is needed for efficient replication restart (Trenz et al., 2006). We therefore examined the functional implications of reduced MLL3 recruitment to forks with p53 defects. Consistently, we find increased MRE11-bound replication sites in WT, but not in p53 null HAP-1 cells when challenged with HU (Figure 3D and figure supplement 1C; 30 MRE11-bound replication sites/cell in WT and 18 in p53 null HAP-1 cells). Collectively, these data suggest a mechanism whereby p53 promotes local chromatin responses that aid MRE11 recruitment to stalled forks, as necessary for replication restart (Trenz et al., 2006).

### p53 suppresses error prone RAD52 at forks

p53 is implicated in suppressing excessive repair by homologous recombination (HR) to balance genomic stability (Bertrand et al., 2004; Saintigny et al., 1999; Sengupta et al., 2004). To further probe the underlying mechanism for genomic instability induced by aberrant p53 S-phase functions, we performed SIRF assays for local RAD51 recruitment to stalled forks as a surrogate marker for HR processes. From previous reports we expected more RAD51 recruitment to forks in the absence of p53 (Bertrand et al., 2004; Gatz and Wiesmuller, 2006). Instead, in p53 null HAP-1 cells, we find less RAD51 at local replication forks (Figure 4A; 24 RAD51-bound replication sites/cell in WT p53 and 19 in p53 null HAP-1 cells). In contrast, HCT116 cells expressing GOF p53 R248W show increased RAD51 fork-localization compared to WT expressing HCT116, as do LOF mutant p53 S47 expressing H1299 cells compared to cells with WT p53 (Figure 4 - figure supplement 1A and 1B). Due to these unexpected differences, we employed p53 null Saos-2 sarcoma cells in comparison to isogenic GOF mutant p53 expressing Saos-2 cells and U2OS cells, which are p53 WT sarcoma cells (Figure 4B). p53 null Saos-2 show an increase in RAD51 SIRF signals, which is repressed with expression of mutant p53 GOF R175H and R273H. Together, these results support no correlation between RAD51 recruitment to forks and fork instability in these cells, but instead suggest alternative causation for the observed genomic instability in p53 mutant cells.

**Figure 4 with 1 supplement.**
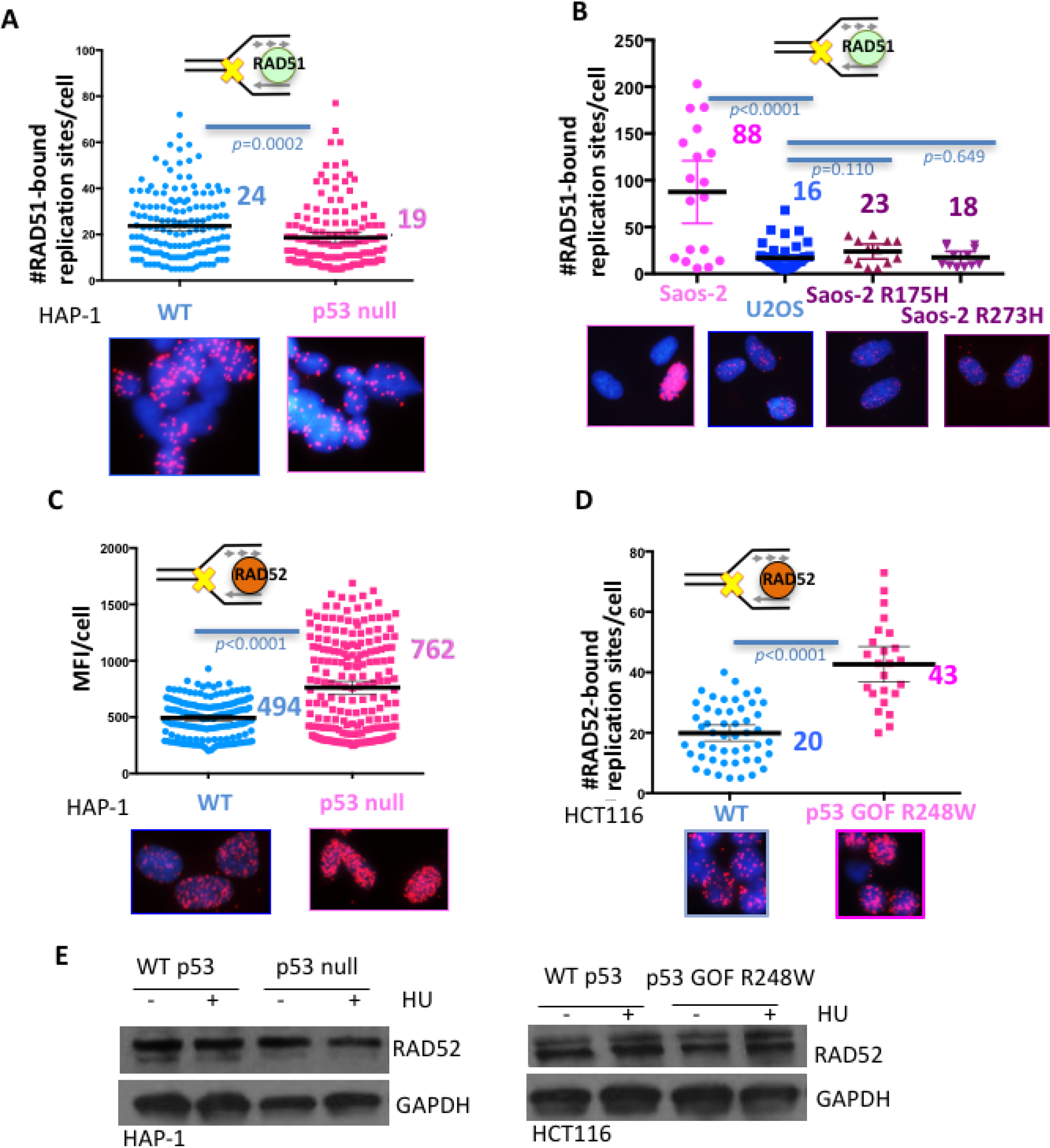
p53 inhibits RAD52 usage at stalled replication forks. (**A**) Quantitation of SIRF assay of RAD51 at HU stalled replication forks in HAP-1 p53 null and WT cells and (**B**) p53 null, R175H and R273H complemented Saos-2 cells, and U2OS cells. (**C**) Quantitation of SIRF assay of RAD52 at HU stalled replication forks in HAP-1 p53 null and WT cells (MFI, mean fluorescence intensity) and (**D**) WT p53 or p53 R248W expressing HCT116 cells. (**E**) Western blot of RAD52 with and without HU (200μM) in p53 null and WT HAP-1 cells, and WT p53 or p53 R248W expressing HCT116 cells. Bars represent the mean and the 95% confidence interval. Significance values are derived from Mann-Whitney analysis.

Defects in p53 do not affect HR repair of induced double-strand breaks, however, they increase spontaneous sister-chromatid exchanges (SCEs) (Willers et al., 2001), which are thought to occur at sites of stalled replication. Reports increasingly suggest that spontaneous SCE is independent of RAD51 and BRCA2 (Bai and Symington, 1996; Clémence Claussin1 and Victor Guryev1, 2017; Ray Chaudhuri et al., 2016), but instead involves the single-strand annealing (SSA) protein RAD52 (Thorpe et al., 2006). We therefore tested whether increased RAD52 recruitment is altered with p53 deletions. Using SIRF analysis, we find a marked increase of RAD52 bound to stalled forks in p53-null cells (Figure 4C). As the signals were too abundant to be individually counted, we used the mean fluorescence intensity (MFI) as a quantitative readout (Figure 4C, MFI of 494 in p53 null and 762 in p53 WT HAP-1 cells). Notably, we see stronger RAD52 recruitment at low compared to high concentration of HU (Figure 4 - figure supplement 1C). The former condition is less favorable for DSB formation, suggesting RAD52 recruitment to stalled forks is stronger than to *bona fide* DNA breaks. Strikingly, we find an increase in RAD52 recruitment in all p53-defective cell lines tested irrespective of the nature of the p53 defect. This includes HCT116 GOF p53 R248W (Figure 4D), Saos-2 p53 null, Saos-2 GOF p53 R175H, Saos-2 GOF p53 R273H, and H1299 LOF p53 S47 cells (Figure 4 - figure supplement 1D and E) compared to respective WT p53 expressing cells. These collective data unexpectedly uncover consistent replication fork pathway tipping towards mutagenic RAD52 processes in p53 defective cells. This pathway imbalance was not caused by transcriptional deregulations in p53 defective cells, as RAD52 protein levels remained unchanged with or without p53, further supporting a transcription-independent function of p53 at stalled forks (Figure 4E). Thus, the observed increased RAD52 recruitment to stalled forks is likely a consequence of defective replication restart.

### p53 suppresses microhomology-mediated end-joining polymerase POLθ

In p53 defective cells, we observed a stark RAD52 recruitment with low HU (Figure 4 - figure supplement 4C), which can lead to reversed replication forks (Neelsen and Lopes, 2015) that provide free ends as substrates for DSB repair pathways. We reasoned that p53 may orchestrate reversed fork outcomes and so protect against aberrant double-strand end recognition by mutagenic DNA end pathways which may include SSA and micro-homology mediated end-joining (MMEJ). POLθ is implicated in promoting error-prone MMEJ at replication-associated DNA ends (Roerink et al., 2014), which may include collapsed or reversed replication forks. We therefore tested if DNA POLθ may contribute to mutagenic events at imbalanced stalled forks. We find an increase of mutant p53 S47 association with POLθ in unchallenged H1299 cells, which is further enhanced with replication stalling (Figure 5D, 24 associations per cell without and 38 with HU). Notably, WT p53-POLθ associations remain limited even with replication stalling (Figure 5C, average of 6 associations without and 15 with HU), suggesting pathway tipping towards mutagenic MMEJ in p53-defective cells.

**Figure 5 with 1 supplement.**
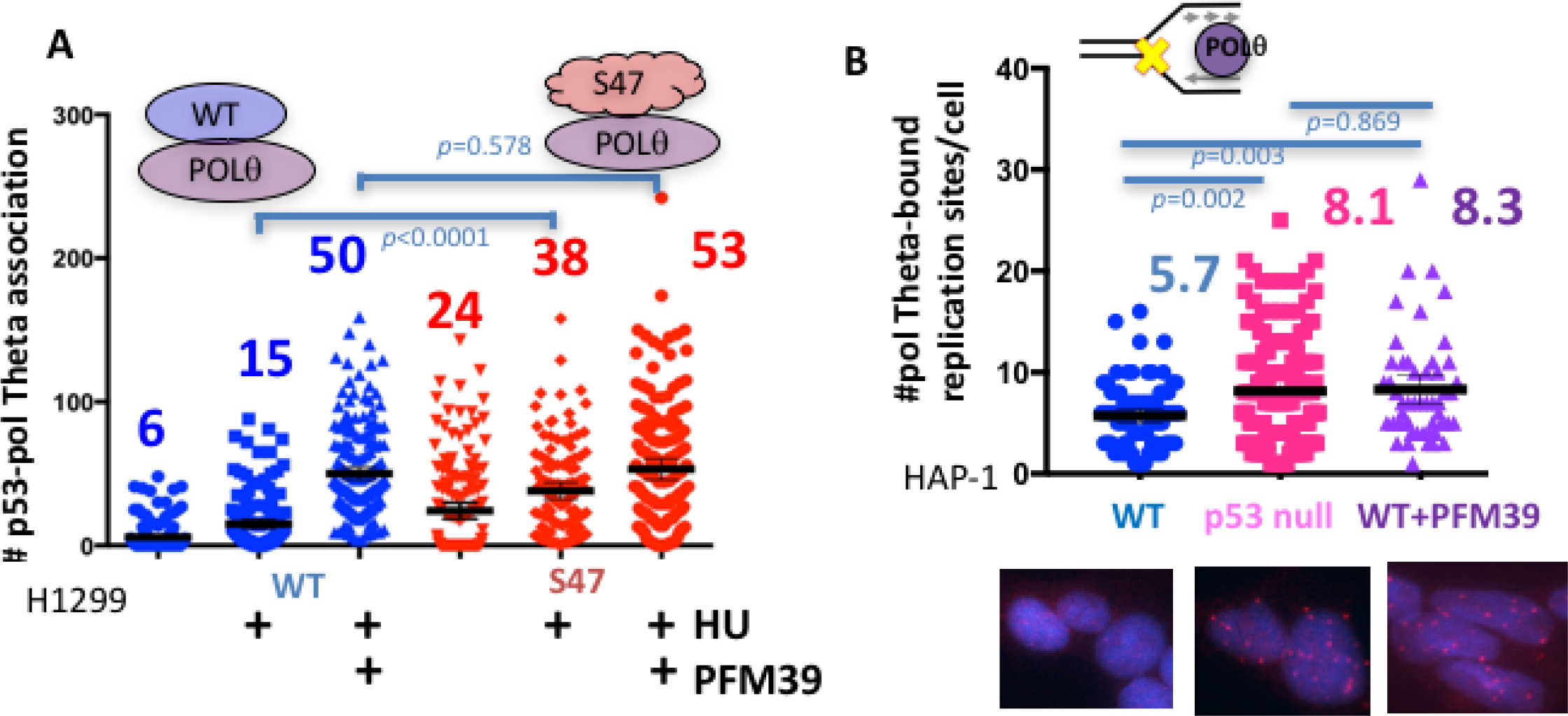
p53 inhibits POLθ usage at stalled replication forks. (**A**) Quantitation of WT p53 or p53 S47 interaction with POLθ by PLA in H1299 cells with or without HU (200μM) and MRE11 inhibitor PFM39 (100μM). (**B**) Quantitation of SIRF assay of POLθ at HU stalled replication forks in HAP-1 p53 null and WT cells. Bars represent the mean and the 95% confidence interval. Significance values are derived from Mann-Whitney analysis.

To test if MRE11-dependent restart is responsible for suppression of error-prone POLθ recruitment, we inactivated the nuclease by inhibition with the specific MRE11 nuclease inhibitor PFM39 (Shibata et al., 2014). Inhibition of MRE11 by PFM39 greatly increased WT p53-POLθ association with replication stalling (Figure 5C, an increase from 15 to 50/cell average with PFM39). This observation suggests that MRE11 inactivation can partially phenocopy p53 deficiency at replication forks. In contrast, p53 S47-POLθ associations were only moderately increased with PFM39 (Figure 5C, increase from 38 to 53 with PFM39), where PFM39 likely blocks residual MRE11 activity in p53 S47 expressing cells. Of note, we find POLθ interactions with RAD52 increased in GOF p53 R248W HCT116 cells (Figure 5 - figure supplement 1A), giving rise to the possibility that POLθ may collaborate with RAD52 in p53 defective cells rather than acting in separate pathways.

To further test pathway imbalance specific to local stalled forks and dependent on p53 status, we performed SIRF against POLθ in HAP-1 cells (Figure 5D). Consistently, we find increased POLθ recruitment to stalled forks in p53-null HAP-1 cells (Figure 5D). Similarly, inactivation of MRE11 nuclease in WT HAP-1 cells causes a significantly increase in recruitment of POLθ to stalled forks. Together these data uncover p53-MRE11 repression of mutagenic RAD52 and POLθ processes at replication forks.

### p53-defective breast cancers show increased mutation signatures typical for RAD52/POLθ

RAD52/SSA and POLθ/MMEJ pathways allow the prediction of specific mutation signatures; SSA predominantly results in larger deletion mutations, while MMEJ is signified by microhomology at repair junctions along with smaller deletions (Jasin and Rothstein, 2013; Wood and Doublie, 2016). We therefore hypothesized that p53 replication-defective cancers may leave a telltale mutagenic pathway signature *in vivo*. We tested this by comparing COSMIC mutational signatures (Stratton et al., 2009) of p53 defective with p53 WT breast cancers reported in the TCGA database (Figure 6, S6A and S6B, Cosine similarity cutoff: 0.617; z-score >1.96). Seven mutational signatures are increased in p53 defective breast cancers (Figure 6B). However, of these seven signatures, only signatures 3 and signature 5 are significantly increased in p53 defective compared to WT p53 breast cancers (Figure 6 - figure supplement 1B); COSMIC signature 3 is defined by larger deletion mutations (>3bp) with microhomology at break junctions, consistent with expected RAD52 and POLθ mutation spectra. COSMIC signature 5 shows T>C transition mutations at ApTpN context with yet unknown etiology. POLθ was reported to have a stark preference for T>C transition mutations (error-rate of 42 ×10^−4^, 4-40 fold higher than any other possible mutation rate) as seen within a known CA<u>T</u>CC hotspot (Arana et al., 2008). Thus, our combined data suggests the possibility of POLθ mediated origin of COSMIC signature 5. Collectively, we find COSMIC signatures are in agreement with increased RAD52 and POLθ pathway usage in p53 mutant breast cancers.

**Figure 6.**
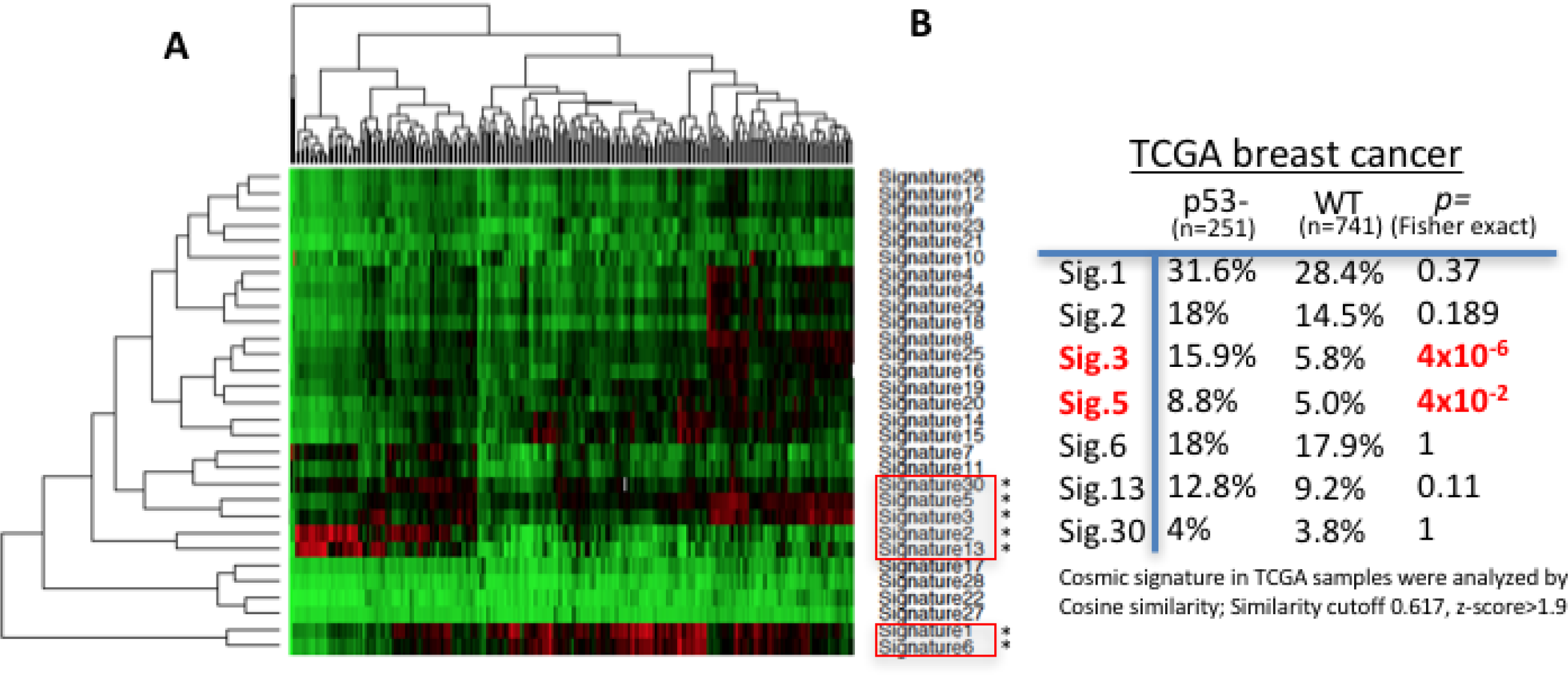
POLθ and RAD52 mutation signatures are upregulated in p53 defective breast cancers. (**A**) Hierarchical clustering of cosmic mutation signatures in p53 defective breast cancer from TCGA database. (**B**) Comparison of frequencies of cosmic signature found unregulated in p53 defective breast cancer with frequencies found in p53 proficient breast cancers.

## DISCUSSION

### p53 suppresses genome instability by orchestrating replication fork homeostasis

Rather than single gene predisposition or select environmental exposure, the strongest drivers for cancer incidence are DNA replication errors. This has been long hypothesized and recently formalized by showing replication errors comprise 2/3 of all mutations in cancers (Tomasetti et al., 2017; Tomasetti and Vogelstein, 2015). This fundamental importance of replication fork maintenance is conserved in bacteria, where stress response and repair proteins primarily protect and stabilize DNA replication forks (Cox et al., 2000).

p53 is the “the guardian of the genome” and the most frequently mutated tumor gene, but its functions in replication genome stability, which is the dominant source of tumor mutations, has been cryptic. The most studied p53 cellular function with regard to tumor-suppression has been its role in transcriptional activation of apoptosis and cell cycle checkpoint. Yet, p53 functions during the DNA damage response linked to genome integrity are transcription activation independent (Bertrand et al., 2004; Janz and Wiesmuller, 2002; Saintigny and Lopez, 2002). Moreover, these classical p53 transactivation activities to promote apoptosis and cell cycle arrest are insufficient to fully explain p53’s role in tumor suppression. This is substantiated by reported p53 separation-of-function mutations, including tumor prone yet greatly transcription activation proficient p53 mutations, such as S47. Conversely, several p53 mutant mice including p53 R175P show that inactivation of apoptosis and senescence by p53 transcription deregulation are insufficient for full inactivation of p53 tumor-suppression functions (Brady et al., 2011; Li et al., 2012; Liu et al., 2004). Taken together these observations point to p53 activities in addition to its transcription activation functions that critically contribute to its tumor suppressor function. Such additional functions may include metabolism and ferroptosis, a new cell death pathway (Li et al., 2012; Zhu et al., 2015).

We here identify a new p53 function in suppressing genome instability at replication forks by promoting MLL3/MRE11-mediated replication pathway homeostasis. Importantly, this activity, which we show is independent of p53 transcription activation roles, avoids mutagenic RAD52/POLθ pathways likely acting at reversed forks (Figure 7). As replication mutations are thought to be the strongest cancer mutation driver and genome instability is associated with tumorigenesis, we propose that the here identified role of p53 as a replication homeostasis keeper to avoid genome instability provides a feasible novel additional p53 tumor suppression function. Moreover, the resulting understanding of p53-mediated genomic stability reconciles previous reports on apoptosis and p53 transactivation-independent roles of p53 for tumor suppression (Phang et al., 2015). So far, the most consistent common defect to both GOF mutant p53 and p53 gene deletion is related to its transcription function in apoptosis and cell cycle arrest. These results revealing a p53 replication-restart function reconcile how GOF and null p53 have different cellular functions and phenotypes, yet can both cause genomic instability implicated for tumor etiology and progression. Supporting this concept, MRE11 impairment, which we show phenocopies p53 defects at stalled forks, promotes progression and invasiveness of mammary hyperplasia in mouse models similar to p53 inactivation (Gupta et al., 2013).

**Figure 7.**
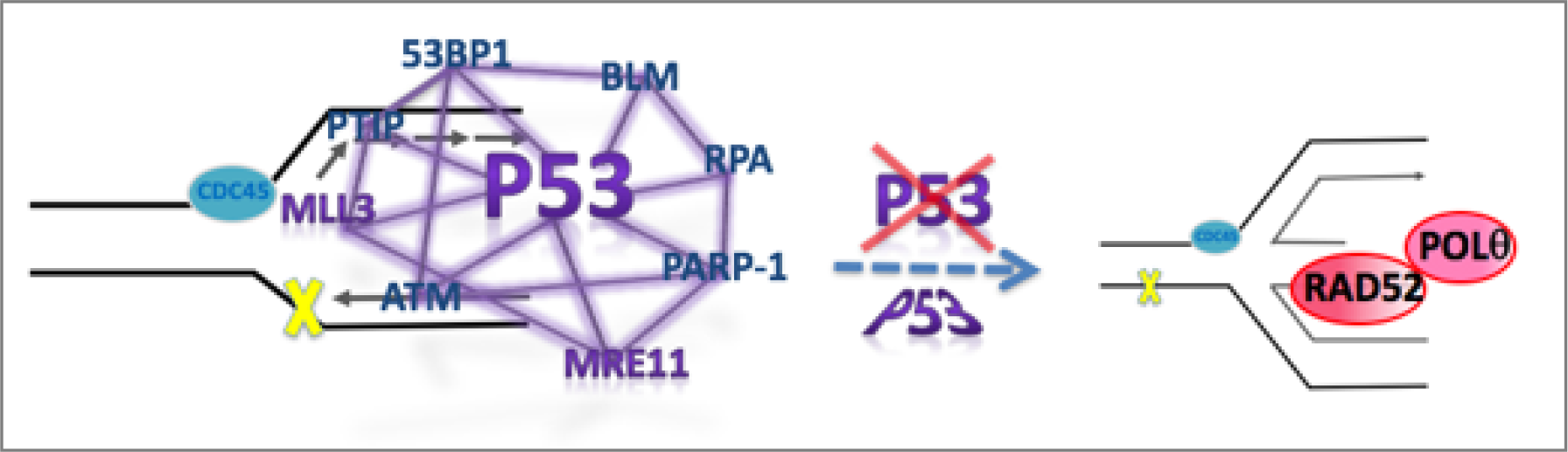
Model for p53-mediated pathway homeostasis. p53 is implicated as a keystone protein that is part of a larger replication restart network. p53 mutations, defects or MRE11 defects tip the replication pathway homeostasis towards increased mutagenic RAD52/POLθ pathways at unprotected stalled forks, such as reversed replication forks, and resulting in deletion and point mutations.

Separation-of-cellular p53 function studies have spanned from apoptosis, cell cycle arrest, epigenetics and stress response to metabolism. Yet, these seemingly separate functions may act together in the context of WT p53 to guard the genome, foremost from genotoxic replication stress. As such, the replication restart function identified here conceptually connects seemingly divergent p53 functions including stress response, genome stability and epigenetics.

We propose that upon activation by replication stress, p53 orchestrates balanced error-free replication restart and suppresses genome instability, which is caused by excessive usage of mutagenic replication pathways when p53 is defective. Importantly, this model implies p53 promotes a replication homeostasis balance at forks for successful proliferation rather than a strict pathway control. If replication stress exceeds a threshold for proper genome maintenance, p53 may dissociate and, as a keystone replication stress regulator, induce cell death, including but not exclusively through apoptosis, as an added safeguard to avoid cellular dysplasia.

### p53 replication fork reactions and implied biological functions

We here find the African-decent tumor variant p53 P47S (S47) to be a separation of function mutation defective in replication restart. p53 is phosphorylated by ATM at S46, which is decreased in p53 S47 (Jennis et al., 2016). Intriguingly, at the adjacent residues D48/D49, p53 can directly interact with single-strand binding protein RPA (Romanova et al., 2004), which is implicated in replication fork remodeling (Neelsen and Lopes, 2015). Specifically, RPA interaction mutations deregulate recombination reactions without affecting transactivation reactions (Romanova et al., 2004). By proximity of these residues and phenotypical commonalities, we suggest that p53 P47S (S47) may also affect RPA interactions. By extension, we propose that ATM phosphorylation of WT p53 may regulate such p53-RPA interactions for the purpose of fork remodeling, as a controlled process for restart balancing. Notably, we find p53 S47 exhibits cellular sensitivity to the DNA cross-linking reagent mitomycin C, which most prominently activates the Fanconi Anemia tumor suppressor pathway. The latest identified Fanconi Anemia tumor suppressor is the RFWD3 ubiquitin ligase that regulates p53 (Feeney et al., 2017; Inano et al., 2017). Furthermore, an Fanconi Anemia phenotypes causing patient mutation in RFWD3 leads to deregulation of RPA reactions at the replication fork (Inano et al., 2017). We therefore propose that p53 could feasibly be a vital player in the Fanconi Anemia pathway through its replication function, and it will be exciting to decipher this feasible relationship.

We establish here that at forks, p53 controls MRE11, a nuclease known to promote restart after replication stalling (Trenz et al., 2006). Other prominent p53 collaborating proteins including BLM helicase (Davies et al., 2007), ATM (Trenz et al., 2006), and PARP-1 (Bryant et al., 2009) all promote replication restart. Our results thus implicate p53 as a potential keystone regulator of a greater restart network at stalled forks involving fork-reversal regulation players (Figure 7). PARP, BLM helicase and RPA promote and repress replication fork reversal (Neelsen and Lopes, 2015). While it is unclear whether fork reversal is required for normal replication restart, it is readily observed in cancer cells at low concentrations of replication stalling agents (Zellweger et al., 2015). In the absence of fork reversal control and stabilization by p53 regulated players, our model suggests that RAD52/POLθ pathways hijack the free DNA end to invade replication ahead or behind the replication fork as an intramolecular reaction. This could in principle lead to deletion and insertion mutations with micro-homology and increased POLθ dependent point mutations. In support of this model, we find COSMIC cancer mutation signature 3 signified by larger deletion mutations with micro-homologies increased in cancers with p53 defects. Additionally we found COSMIC signature 5 to be increased in p53 defective breast cancers, which shows T>C transition mutations at ApTpN context with so far unknown etiology. However, our analysis shows that this signature is consistent with POLθ-mediated mutations: POLθ shows a striking preference for T>C transition mutations (error-rate of 42×10^−4^, 4-40 fold higher than any other possible mutation rate) as seen within a known CA<u>T</u>CC hotspot (Arana et al., 2008). Based on our data, we therefore suggest that POLθ mediated mutagenesis may contribute to COSMIC signature 5 etiology.

### Implications of p53 restart function in therapy resistance

In mammary tumor suppression, p53 cooperates with BRCA1/2 (Jonkers et al., 2001; Ludwig et al., 1997), which is often associated with more aggressive and resistant tumors, although the mechanism of this collaboration has been elusive. Therapy resistance in BRCA defective cells can arise through secondary mutations. Specifically, gene internal deletion and/or point mutations within the BRCA2 gene can restore reading frames of BRCA2 mutated stop codons in CAPAN-1 pancreatic and POE ovarian cancer cells (Sakai et al., 2009; Sakai et al., 2008). Interestingly, while BRCA2 peptide expression is restored, some of these new BRCA2 peptides conferring resistance have extensive internal deletion mutations. Such deletion mutation etiology is consistent with both RAD52 and POLθ pathways providing a specific and testable mechanism for development of resistance.

Our model of RAD52/ POLθ pathway increase at stalled replication forks promoting secondary mutation driving resistance is further supported by our understanding of tumor resistance biology. Triple negative breast cancers as well as serous ovarian cancers, which almost exclusively harbor p53 mutations, are often initially sensitive to therapy such as cis-platin drugs (Luvero et al., 2014; Wahba and El-Hadaad, 2015), which cause replication stalling. After multiple treatments and opportunities for RAD52/POLθ mediated mutagenic events at stalled forks, secondary mutations can become fixed and in turn promote survival and resistance. In this scenario, mutation fixation is not necessary to promote a proliferative advantage *per se*. Rather it could arise from stochastic and opportunistic replication stalling events promoted by the therapeutic drug dependent on the replication program of the tissue type, consistent with both a neutral mutation evolution theory (Sottoriva et al., 2015) and replication errors driving tumor etiology (Tomasetti et al., 2017).

We here identified a new p53 role for suppressing genome instability by orchestrating balanced replication fork homeostasis. Importantly, this role is derailed in both p53 null and GOF p53 mutants, which is the only loss-of-function ascribed to both aside from transcriptional deregulation of apoptosis and cell cycle checkpoint. Our observations and concepts reconcile prevailing paradoxes of divergent p53 functions. They furthermore imply specific changes in strategies for cancer patient care: our model suggests that inhibition of RAD52/POLθ pathways as adjuvant therapy concomitant with initial conventional therapy could offer an actionable strategy for ameliorating aggressive tumor evolution and secondary mutations leading to resistance in p53 defective tumors. The finding that p53 is a key-protein in error-free replication restart may explain why p53 mutations are a dominant cause of cancer genome instability.

## Acknowledgments

We thank Dr. Guillermina Lozano (UT MD Anderson) for discussion and reagents (MEF p53 R172H and p53 R172P, and Saos-2 null, R175H and R273H cells). We thank Dr. Ron DePinho and Dr. John Tainer (UT MD Anderson) for critical reading of the manuscript. PFM39 was synthesized by MD Anderson Cancer Center pharmaceutical chemistry core facility. This work was supported by the National Cancer Institute of the National Institutes of Health under Award K22CA175262 and by CPRIT Award R1312. K.S. is a Rita Allen Foundation Fellow, a CPRIT Scholar in Cancer Biology and an Andrew Sabin Family Foundation Fellow.

## Contribution

S.R.; experimental design, data curation, data analysis, manuscript editing. K.T; data curation, data analysis, manuscript editing. S. P., data curation, data analysis. J.W.L. Manuscript editing, J.L., data analysis. M.M. resources, critical discussion. K.S.; conceptualization, experimental design, data curation, data analysis, supervision, manuscript writing and editing, figure design, resources, funding acquisition

## Competing interests

There are no competing interests.

## MATERIAL AND METHODS

### Cell lines and reagents

HAP-1 parental and HAP-1 TP53 null (Horizon Discovery) cells were grown in Iscove’s modified Dulbecco’s medium (Life Technologies) supplemented with 10% fetal bovine serum (Gemini Bio products) and 100units/ml Pen-Strep (Life Technologies). H1299 small lung cell carcinoma cells expressing doxycycline inducible human WT and S47 mutant p53 constructs were previously described (Jennis et al., 2016) and grown in Dulbecco’s modified Eagle medium supplemented with 10% fetal bovine serum and 100 units/ml Pen-Strep. P53 protein expression was induced by 0.5μg/ml Doxycycline (Sigma-Aldrich). MCF10A p53 null cells were obtained from Thermo Fisher Scientific and grown in DMEM/F12 medium supplemented with 5% horse serum (Gibco), 20ng/ml EGF (Thermo Fisher Scientific), 0.5 μg/ml hydrochortisone (Sigma-Aldrich), 100ng/ml Cholera toxin (Sigma-Aldrich), 10 μg/ml Insulin (Sigma-Aldrich), 5 mM Hepes (Gibco), 100 units/ml Pen-Strep. MEF harboring p53 mutations R172P, R172H were previously described (Liu et al., 2004), and MEF harboring p53 mutations S47 and WT p53 and p53 null MEF were previously described (Jennis et al., 2016), and obtained from the Guillermina Lozano lab and the Maureen Murphy lab, respectively. MEFs were grown in Dulbecco’s modified Eagle medium supplemented with 10% fetal bovine serum and 100 units/ml Pen-Strep and 2mM glutamine. MEFs were generated from C57BL/6J mice with mixed sex background. HCT116 parental and CRISPR engineered mutant cells (R248W/-) were obtained from Thermo Fisher Scientific and grown in McCoy’s 5a media (Lonza) with 10% fetal bovine serum and 100 units/ml Pen-Strep. Saos-2 cells complemented with gain of function p53 mutants were previously described (Xiong et al., 2014), provided by Dr. Guillermina Lozano’s lab and maintained in DMEM (Life Technologies) with 10% fetal bovine serum and 100 units/ml Pen-Strep. Cell lines have been authenticated by short tandem repeat (STR) profile analysis and genotyping. All cells were grown at 37°C and 5% CO_2_.

### DNA Fiber assay

DNA fiber spreading experiments were performed as previously described (Schlacher et al., 2011). Briefly, cells were pulsed with EdU (5-125 μM), CldU (50 μM) or IdU (50 μM), washed with PBS, and then incubated with hydroxyurea (200-400 μM) and CldU (50 μM) for 4-5 hours as indicated. The cells were harvested, resuspended in PBS and lysed on a microscope slide with lysis buffer (20 mM Tris-Cl, 50 mM SDS, 100 mM EDTA). DNA was allowed to attach for 5.5 minutes before spreading by gravity. Slides were fixed in methanol/acetic acid (3:1), before DNA denaturation with 2.5 N HCl and neutralization with PBS (pH 8, and subsequent pH 7.5 washes). Slides were blocked with 10% goat serum and 0.1% Triton X in PBS. IdU/CldU fibers were stained using standard immunostaining with antibodies against IdU (BrdU, Beckton Dickinson, 1:100) and CldU (BrdU, Novus Biological, 1:200) was performed before mounting slides with Prolong Gold (Invitrogen). IdU/CldU Fibers were imaged using a Nikon Eclipse Ti-U inverted microscope and analyzed using ImageJ software. Between 90-320 fibers were scored per experiment and number of stalled forks was calculated as the number of IdU tracts (green only) divided by the number of IdU tracts plus the number of IdU-CldU tracts (green followed by red). The number of newly initiated forks was calculated as the number of CldU tracts (red only) divided by the number of IdU tracts plus the number of IdU-CldU tracts (green followed by red) plus the number of CldU tracts (red only).

### SIRF assay

Cells were pulse treated with EdU, washed 2 times with PBS and subsequently treated with HU (0.2 μM) for 4 hours. Cells were fixed, permeabilized with 0.25% TritonX, and a click-iT reaction was performed using biotin azide (Life Technologies) according to manufacturer’s instructions. After incubation with primary antibodies, a Duolink proximity ligation assay (Sigma-Aldrich) was performed with mouse/rabbit detection red reagents according to the manufacturer’s instructions. Slides were stained with DAPI and mounted with Prolong Gold before imaging using Nikon Eclipse Ti-U inverted microscope. Signals were analyzed using Duolink software, ImageJ and Nikon NIS elements, in addition to hand-counting of PLA signals. Data of repeated experiments was combined, and statistical analysis was performed using Prism6 software.

### Proximity Ligation Assays

H1299 cells were treated with 0.5 μg/ml doxycycline (Sigma-Aldrich) for 48 hours to induce expression of WT and mutant p53 and subsequently treated with 100 μM PFM39 (synthesized by the MD Anderson Cancer Center pharmaceutical chemistry core facility according to (Shibata et al., 2014)) for 30 minutes, followed by 0.2 mM HU for 4 hours, as indicated. Cells were fixed, permeabilized and blocked as described above and incubated with antibodies against p53 and POLθ as indicated. Finally, a Duolink PLA (Sigma-Aldrich) was performed according to manufacturer’s instructions. Slides were stained with DAPI and mounted with Prolong Gold before imaging using Nikon Eclipse Ti-U inverted microscope. Signals were analyzed using Duolink software, ImageJ and hand-counted. Data of repeated experiments was combined, and statistical analysis was performed using Prism6 software.

### Immunoblotting and antibodies

For Western blots, cells were treated with 0.3 mM HU for 4 hours, harvested and directly lysed in Laemmli buffer (Bio-Rad), boiled for 5 minutes and loaded on SDS-PAGE gels. Antibodies used for immunoblots in SIRF and PLA are as follows: MLL3 (Abcam 1:100), MRE11 (Abcam 12D7 1:200), RAD52 (Santa Cruz F7 1:50), POL θ (Abcam 1:100), RAD51 (Abcam 14B4 1:200), mouse biotin (Sigma-Aldrich BN-34 1:100) and rabbit biotin (Cell Signaling D5A7 1:200).

### Genomic instability assay

Cells were incubated with 50 μM EdU for one hour and subsequently with 0.2 mM HU and 2 μg/ml cytochalasin B (Sigma-Aldrich) for 5 hours. Cells were then collected, washed and treated with cytochalasin B for 20 hours to further capture arrested cells after division that previously were EdU labeled. Post incubation, cells were harvested and spun onto slides using a cytospin for 3 minutes at low acceleration setting. Cells were then fixed, permeabilized and click-iT reaction was performed with Alexa fluor 488 azide according to manufacturer’s instructions. Slides stained with DAPI and mounted with Prolong Gold before imaging using Nikon Eclipse Ti-U inverted microscope. EdU positive cells and micronuclei were scored manually and using ImageJ software. Prism was used for statistical analysis of combined repeat experiments.

### Cell Survival Assays

Cell viability was determined using the colorimetric MTS assay. Cells (1-2 × 10^3^ cells) were seeded into 96-well plates for 24 h and then exposed to varying concentrations of HU or MMC (Sigma-Aldrich) as indicated. After untreated control cells obtained ~80% confluence, the MTS assay was performed according to manufacturer’s instructions (CellTiter 96 AQueous One Solution Cell Proliferation Assay, Promega). Experiments were performed in quadruplicate and repeated independently. Data was analyzed using Prism6 software and represents the mean +/− Standard error of the mean (SEM).

### TCGA Computational Analysis

The mutation annotation file (MAF) for 992 samples was downloaded from BROAD TCGA GDAC website (http://firebrowse.org/?cohort=BRCA&downloaddialog=true). The mutation spectrum of each sample was estimated by calculating the fraction of 96 possible mutation substitutions defined in (Alexandrov et al., 2013) The cosine similarity score is computed for all pair-wise combinations of mutation spectrum of samples and 31 cosmic mutation signatures (http://cancer.sanger.ac.uk/cosmic/signatures). Z-score is calculated based on the distribution of all cosine similarity score 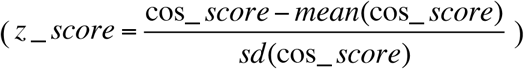. A z score greater than 1. 96 indicates the sample could contain the corresponding cosmic signature.

### Statistical Analysis

For SIRF assays, PLA signals were analyzed using Duolink Image Tool software and Nikon NIS elements software. A total of 50-300 nuclei were counted for each experimental condition. Data represents pooled experiments of 2-4 experiments. Mann-Whitney statistical test for significance was performed using GraphPad Prism version 6 and is indicated in the respective figures and figure legends. For DNA fiber assays, between 90-300 fibers were analyzed using ImageJ software. Unpaired Student t-test was performed using GraphPad Prism version 6 as indicated in the figures and figure legends. For genomic instability were analyzed using NIS elements software. Unpaired Student t-test was performed using GraphPad Prism version 6 to determine p value results as indicated in the figures and figure legends. For TCGA Computational Analaysis, Fisher Exact Test was calculated using GraphPad Prism software.

---

**Figure 1 – figure supplement 1.**
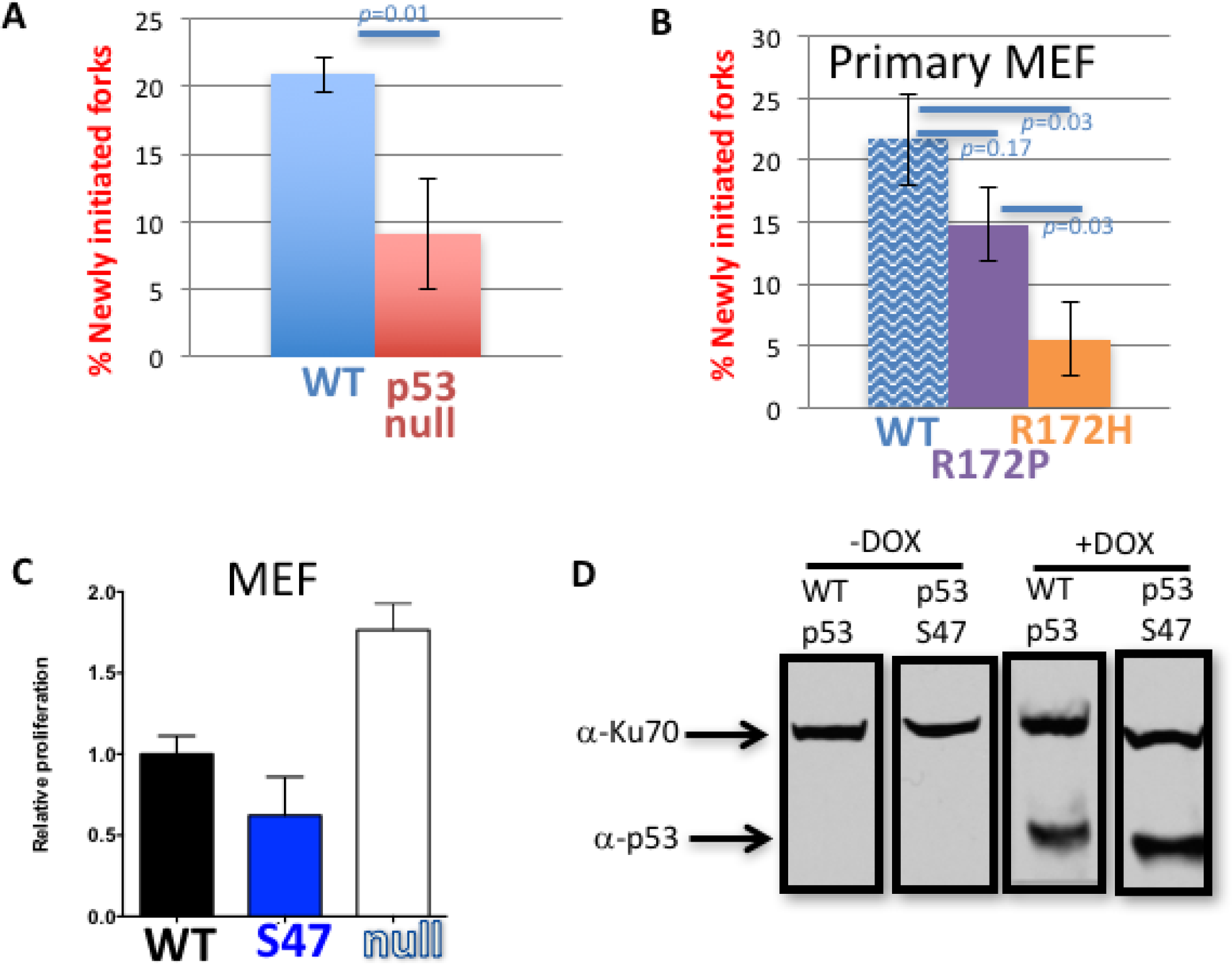
**(A)** DNA fibers analysis of newly initiated forks of p53 null and WT HAP-1 cells with replication stalling (200μM). **(B)** DNA fibers analysis of newly initiated forks of primary MEF with WT, GOF p53 R172P and p53 R17ZH cells with replication stalling (300μM). **(C)** Expression of p53 S47 and p53 WT suppress proliferation compared to primary MET without p53 as measured by MTT proliferation assay, consistent with intact cell cycle checkpoint functions in the mutant. **(D)**Doxycyclire induced expression of WT and mutant p53 S47 in H1299 cells.

**Figure 2 – figure supplement 2.**
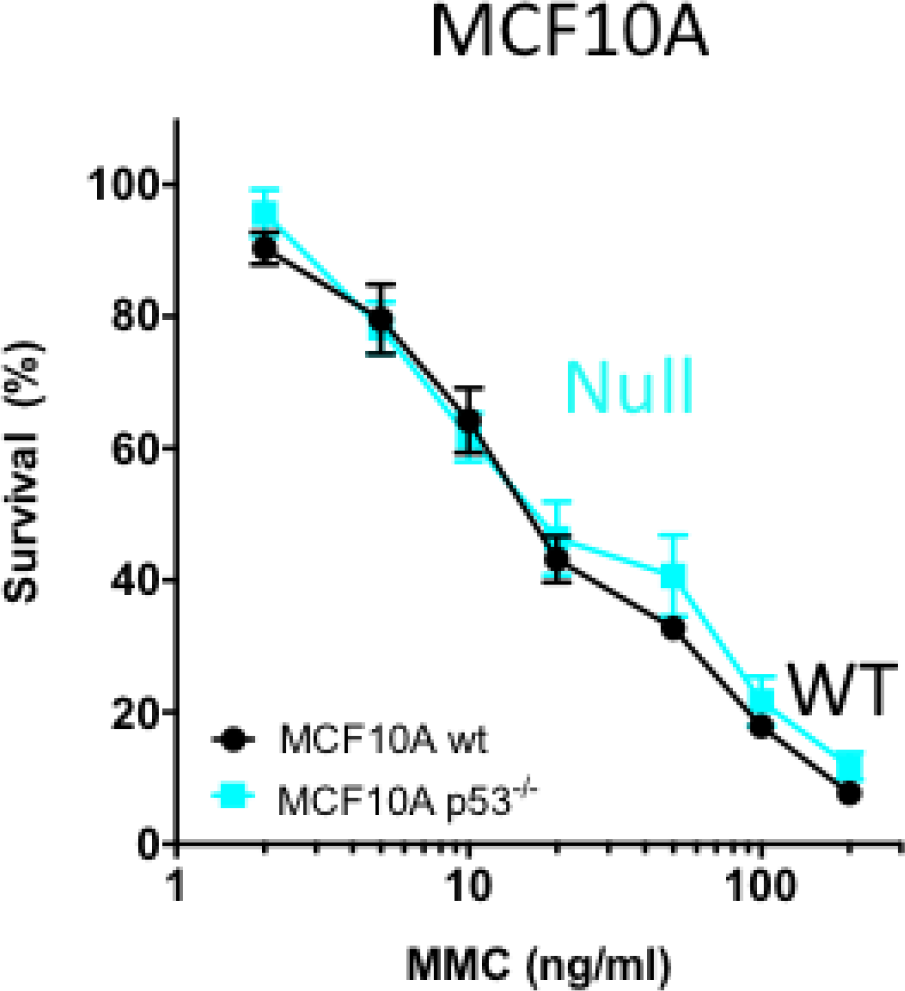
p53 null MCF10A cells show similar sensitivity to mitomycin C compared to WT p53 MCF10A cells.

**Figure 3 – figure supplement 3.**
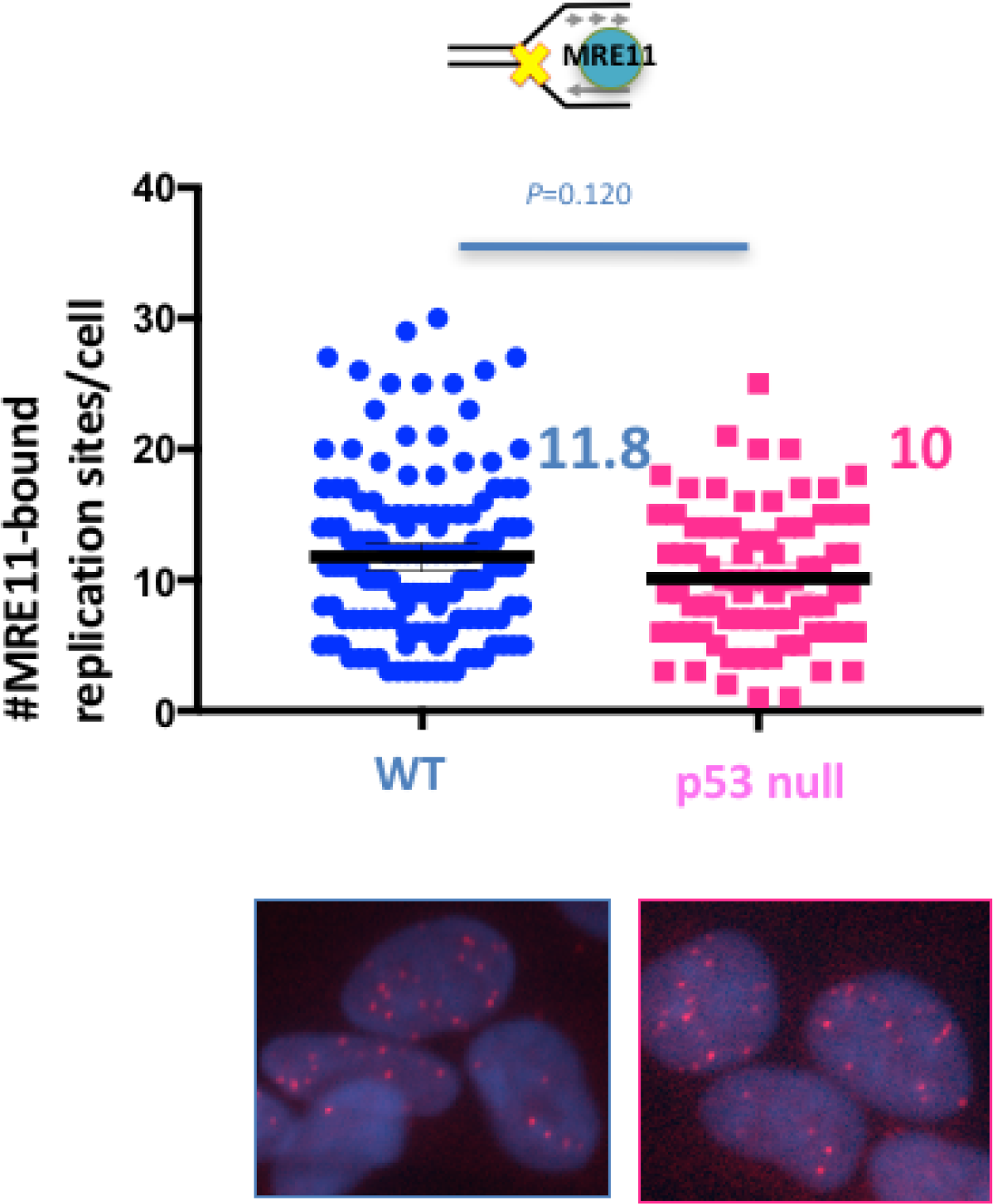
Quantitation of SIRF assay of MRE11 in unperturbed HAP-1 p53 null and WT cells. Bars represent the mean and the 95% confidence interval, Significance values are derived from Mann-Whitney analysis.

**Figure 4 – figure supplement 4.**
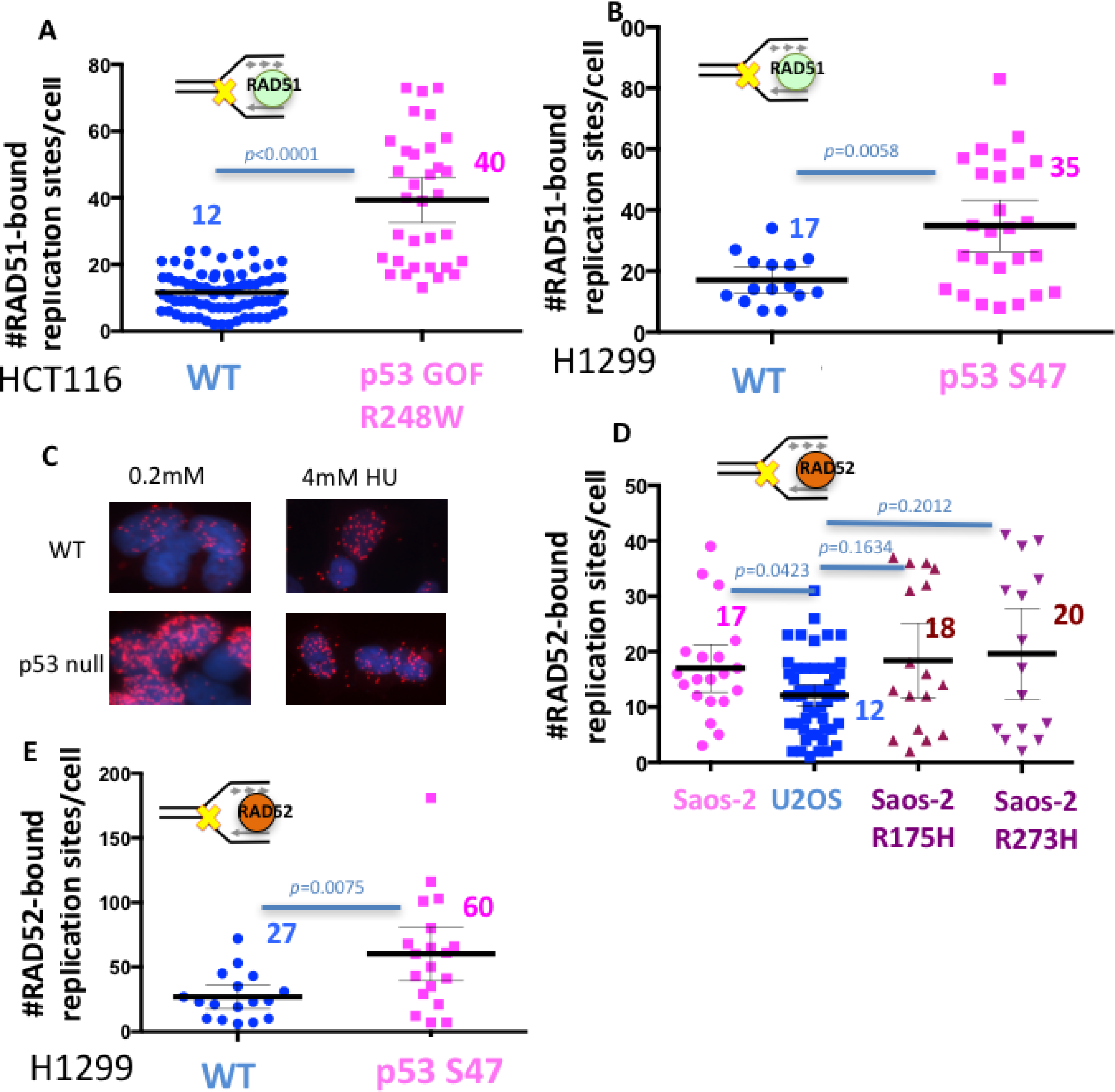
**(A)** Quantitation of SIRF assay of RAD51 at HU stalled replication forks in pS3 R248W and WT p53 HCT116 cells. **(B)** Quantitation of SIRF assay of RAD51 at HU stalled replication forks in p53 547 and WT p53 H1299 cells. **(C)** Representative images of SIRF assay of RADS2 in HAP-1 cells at low (O.ZmM) and high (4mM) HU concentrations show preferential binding of RADS2 at stalled forks with low HU concentrations compared to high HU concentrations, which are more favorable for break formation. **(D)** Quantitation of SIRF assay of RADS2 at HU stalled replication forks in p53 R17SH, p53 R273H and p53 null Saos-2 sarcoma cells, and WT p53 U205 sarcoma cells. **(E)** Quantitation of SIRF assay of RAD52 at HU stalled replication forks in pS3 S47 and WT pS3 H1299 cells. Bars represent the mean and the 95% confidence interval. Significance values are derived from Mann-Whitney analysis.

**Figure 5 – figure supplement 5.**
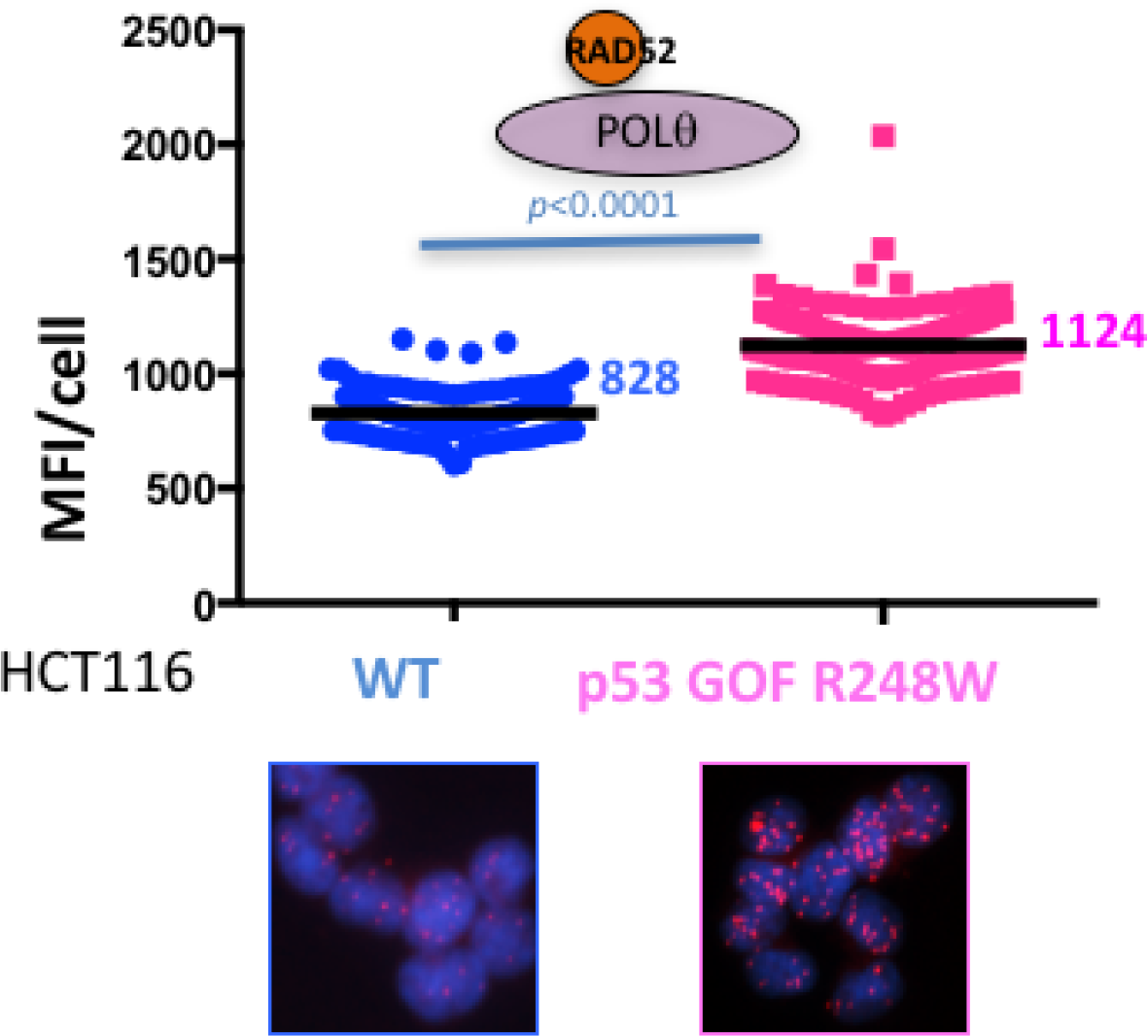
PLA assay between RAD52 and POLθ in p53 R248W and WT p53 HCT116 cells, Bars represent the and the 95% confidence interval. Significance values are derived from Mann-Whitney analysis.

**Figure 6 – figure supplement 6.**
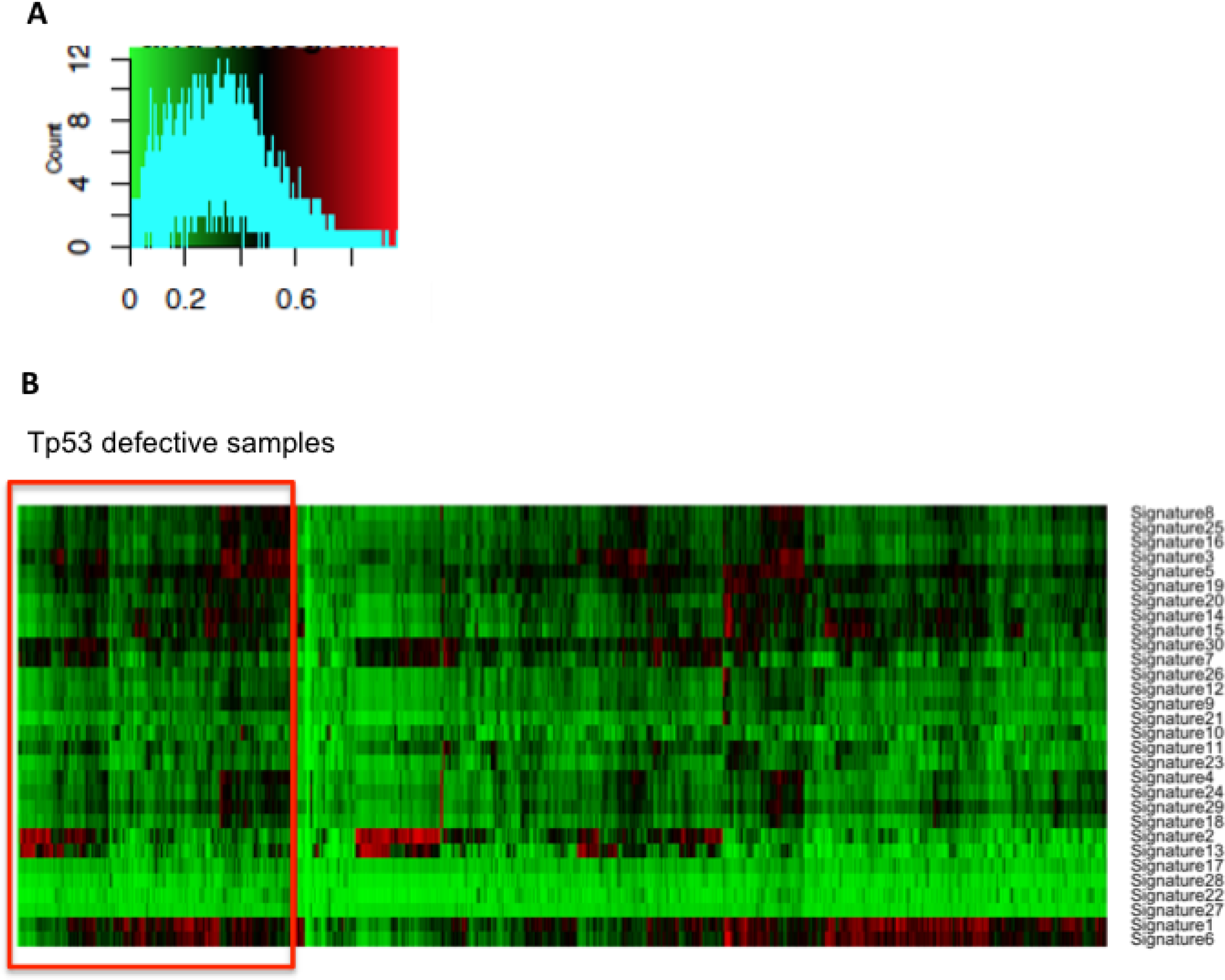
**(A)** Color key histogram showing the distribution of the similarity score **(B)** Hierarchical clustering of cosmic mutation signatures in p53 defective and proficient breast cancer from TCGA database.

